# Goal-directed action preparation in humans entails a mixture of corticospinal neural computations

**DOI:** 10.1101/2024.07.08.602530

**Authors:** Corey G. Wadsley, Thuan Nguyen, Chris Horton, Ian Greenhouse

**Author notes:** Corresponding author: Ian Greenhouse Department of Human Physiology 181 Esslinger Hall, 1525 University St. Eugene, OR 97403.

## Abstract

The seemingly effortless ability of humans to transition from thinking about actions to initiating them relies on sculpting corticospinal output from primary motor cortex. This study tested whether canonical additive and multiplicative neural computations, well-described in sensory systems, generalize to the corticospinal pathway during human action preparation. We used non-invasive brain stimulation to measure corticospinal input-output across varying action preparation contexts during instructed-delay finger response tasks. Goal-directed action preparation was marked by increased multiplicative gain of corticospinal projections to task-relevant muscles and additive suppression of corticospinal projections to non-selected and task-irrelevant muscles. Individuals who modulated corticospinal gain to a greater extent were faster to initiate prepared responses. Our findings provide physiological evidence of combined additive suppression and gain modulation in the human motor system. We propose these computations support action preparation by enhancing the contrast between selected motor representations and surrounding background activity to facilitate response selection and execution.

Abstract figure legend.
Goal-directed action preparation shapes corticospinal output across selected, nonselected, and task-irrelevant motor representations. This study examined whether additive and multiplicative neural computations, common in sensory systems, occur within the corticospinal pathway during action preparation. We probed corticospinal input-output during the performance of various instructed-delay response tasks by applying a range of transcranial magnetic stimulation (TMS) intensities (input) over the primary motor cortex and measuring the resultant motor-evoked potentials (output) from the hand. We found that goal-directed action preparation increases corticospinal gain multiplicatively in task-relevant motor representations while additively suppressing nonselected and irrelevant representations. Greater gain modulation predicted faster responses, highlighting how these computations can enhance signal-to-noise (SNR) to enable efficient action selection and execution in the human motor system.

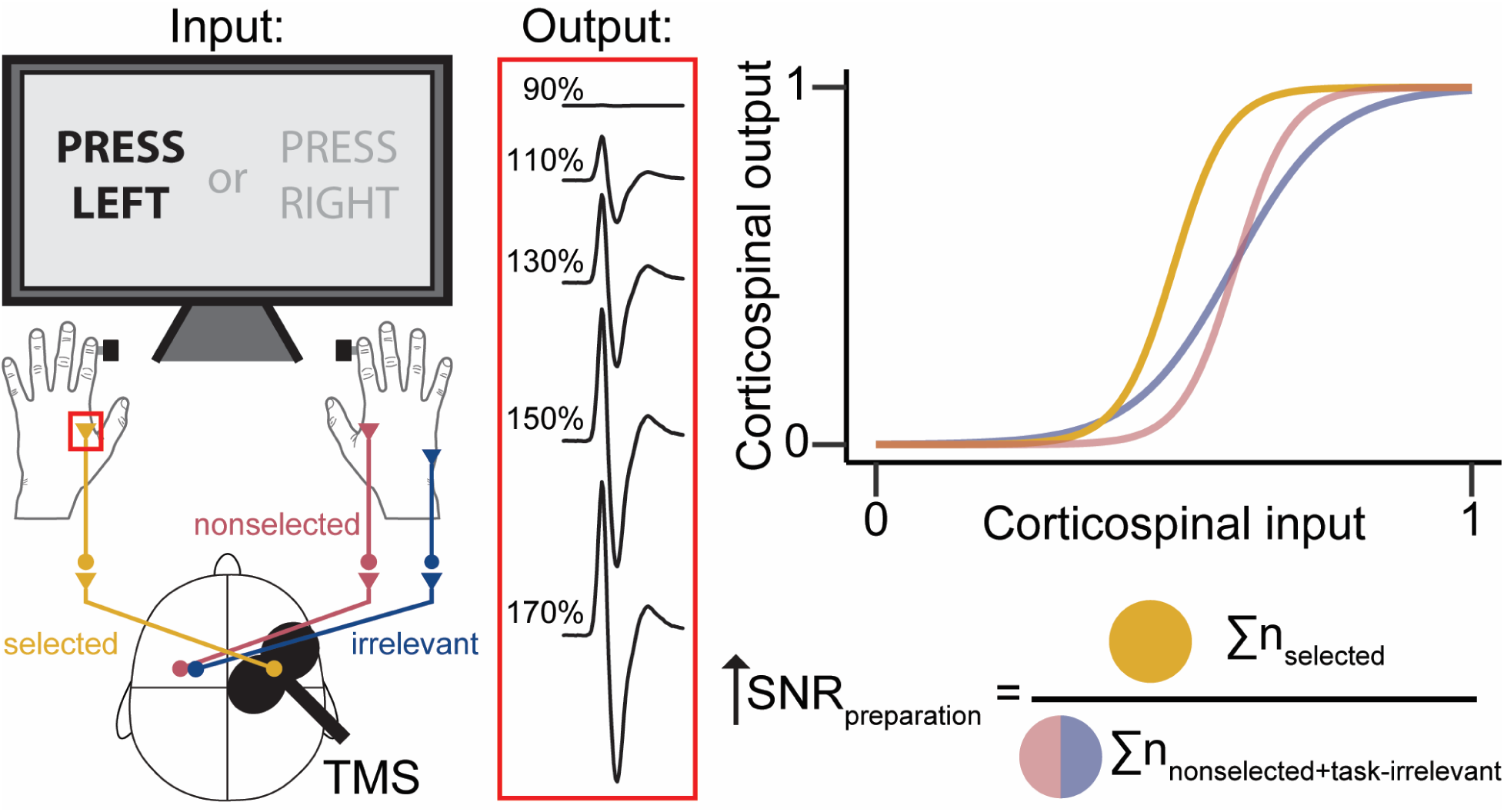

**Key points:**

- Neural computations determine what information is transmitted through brain circuits.
- We investigated whether the motor system uses computations similar to those observed in sensory systems by noninvasively stimulating the corticospinal pathway in humans during goal-directed action preparation.
- We discovered physiological evidence that corticospinal projections to behaviorally relevant muscles exhibit nonlinear gain computations, while projections to behaviorally irrelevant muscles exhibit linear suppression.
- Our findings suggest that certain computational principles generalize to the human motor system and serve to enhance the contrast between relevant and background neural activity.
- These results indicate that neural computations during goal-directed action preparation may support motor control by increasing signal-to-noise within the corticospinal pathway.

## Introduction

Humans are experts at using their fingers for goal-directed actions to interact with their environment effectively (Xu *et al*., 2024). This behavioral repertoire is supported by a specially evolved neural architecture comprising excitatory and inhibitory mechanisms that sculpt corticospinal (CS) pathway output during goal-directed action preparation (Prut & Fetz, 1999; Duque *et al*., 2017; Bundt & Huster, 2024). The anatomy of the distributed cortical and spinal circuits that modulate CS output during action preparation and execution is well characterized (Wang *et al*., 2017; Derosiere & Duque, 2020). However, the precise neural computations that unfold throughout the transition from action preparation to execution are not (Churchland & Shenoy, 2024). Candidate computations have been proposed but are based on neuronal circuit modeling and experimental evidence from investigations of non-motor neural systems and behaviors (Greenhouse, 2022). Thus, it is unclear whether computational principles from non-motor circuits generalize to the human motor system and account for the top-down modulation of CS output during action preparation.

Influential models of goal-directed action preparation postulate that inhibition of the CS pathway acts as a functional gate, with the removal of inhibition allowing selective excitatory signals to pass through to the musculature and produce intended actions (Zagha *et al*., 2015; Soteropoulos, 2018). Yet, physiological recordings from non-human primates have provided mixed evidence for gating mechanisms within the CS pathway (Merchant *et al*., 2008; Kaufman *et al*., 2010, 2013). Indeed, (dis)inhibition supports a variety of computations other than gating in non-motor neural circuits (Katzner *et al*., 2011; Lee *et al*., 2012; Mischler *et al*., 2023). Multifaceted computations are also likely required in the motor system since movements are not static but instead defined by various kinetic and kinematic features that characterize a given action. Therefore, alternative interpretations to the functional gating account should be considered to better explain how preparatory activity determines CS pathway excitability to shape ensuing actions.

The selection and timing of goal-directed actions may be determined by neural computations similar to, or bound to, those of non-motor circuits (Beste *et al*., 2023). Such computations, responsible for sculpting descending signals, are expected to manifest in distinct patterns of CS input-output during goal-directed action preparation. One candidate computation is gain modulation, defined in studies of sensory processing as a multiplicative (non-linear) change in a system‘s input-output relationship and a proposed feature of canonical normalization processes within the brain (Hillyard *et al*., 1998; Salinas & Thier, 2000; Pi *et al*., 2013). Computational models indicate gain modulation is important in determining primary motor cortex (M1) output (Pouget & Snyder, 2000; Stroud *et al*., 2018), the predominant source of descending signals of the human CS tract (Welniarz *et al*., 2017). Additive modulation, defined by a linear shift across an input-output spectrum (Silver, 2010), is an alternative candidate computation that may account for CS input-output adjustments during action preparation. Computational models of additive modulation indicate effective changes in an input-output threshold (Mitchell & Silver, 2003), whereby widespread suppression can stabilize a neuronal population by reducing the preponderance of spontaneous firing (Ferguson & Cardin, 2020). During action preparation, additive modulation may function as the ‘quiet before the storm’ to prepare the CS pathway for impending descending commands.

A third alternative is that multiplicative and additive computations are concurrent and complementary. A precedent for synergistic computations can be found in the vision system. Extending a divisive normalization framework (Carandini & Heeger, 2012) to the motor system, the activity of a motor representation selected for action can be divided by the summed activity of alternative non-selected and bystander motor representations, producing enhanced signal-to-noise (Georgopoulos & Stefanis, 2007; Greenhouse, 2022). A combination of multiplicative and additive computations could serve to separate figure from ground in a manner resembling neural sensory processing and would be consistent with a general neurocomputational principle throughout the human nervous system akin to exponentiation, linear filtering, and divisive normalization (Carandini & Heeger, 2012).

Evidence from transcranial magnetic stimulation (TMS) studies indicates that the motor system engages modulatory processes analogous to those of sensory systems. TMS is a noninvasive brain stimulation technique used to elicit motor-evoked potentials (MEPs) as an index of CS output (Bestmann & Duque, 2016). Seminal studies have demonstrated surround inhibition of MEPs within the CS pathway, whereby selected motor representations are potentiated while surround representations are suppressed during action initiation (Stinear & Byblow, 2003; Sohn & Hallett, 2004). Such surround inhibition is thought to ‘sharpen’ motor representations (Georgopoulos & Stefanis, 2007), a motif that resembles surround suppression observed consistently in the visual (Allman *et al*., 1985; Angelucci *et al*., 2017), auditory (Wang *et al*., 2012; Gilday *et al*., 2023), somatosensory (Foeller *et al*., 2005), and olfactory (Aungst *et al*., 2003) systems. Other TMS studies have identified distinct modulation of MEPs via intra-hemispheric and inter-hemispheric mechanisms across various action preparation contexts (e.g., Derosiere & Duque, 2020). However, these studies have typically used a fixed intensity and could not determine whether MEP modulation is part of additive or multiplicative computations across the entire CS input-output spectrum. Therefore, it remains unclear whether the human CS pathway performs canonical additive and multiplicative computations during action preparation.

In this study, we tested the presence of multiplicative and additive computations in human CS input-output during the behavioral state change from rest to action preparation of goal-directed finger movements (Figure 1a). We hypothesized that goal-directed action preparation involves a concurrent gain increase and additive suppression to enhance the contrast between selected and background motor representations along the CS pathway. To test this hypothesis, we used TMS to noninvasively probe the CS input-output of a target hand motor representation during instructed-delay reaction time tasks. These tasks involved either a choice between two fingers within one hand or between the two hands making it possible to evaluate CS input-output across various contexts, including when the same stimulated motor representation was selected, nonselected but task-relevant, or entirely task-irrelevant to action preparation. This design enabled us to characterize computations within the same CS pathway under different behavioral contexts.

**Figure 1:**
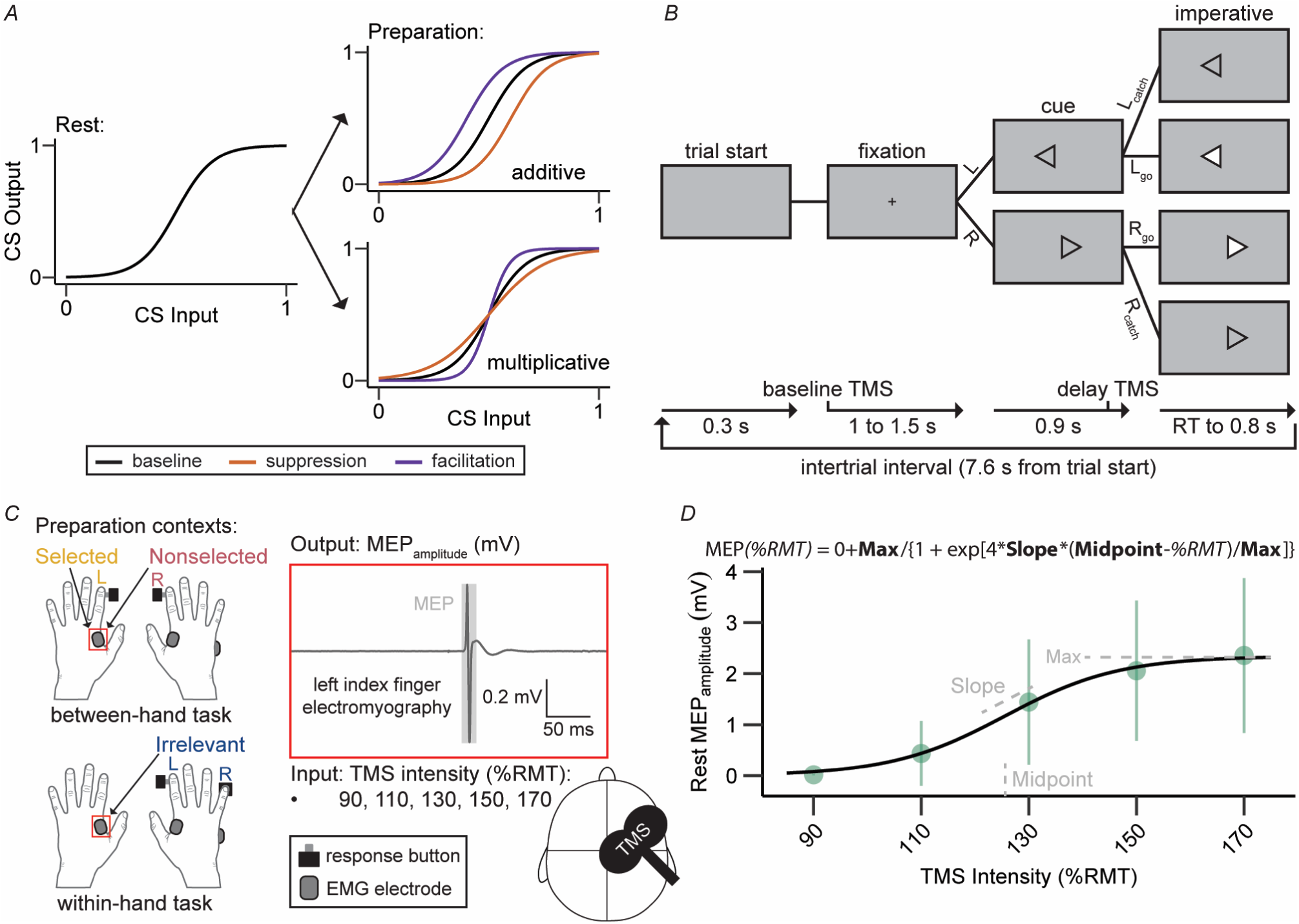
Assessment of human CS input-output during goal-directed action preparation. **a**, Corticospinal (CS) input-output can exhibit additive and/or multiplicative modulation from baseline rest to a preparatory state. **b**, Goal-directed action preparation was assessed with an instructed-delay two-choice reaction time task requiring left (L) or right (R) responses. Most trials were go trials during which the imperative stimulus appeared until a reaction time (RT) was recorded or 0.8 s elapsed. A subset of catch trials (∼8%) was included to prevent participants from predicting imperative stimulus onset. **c**, The experiment was split into between-hand and within-hand tasks, enabling the assessment of CS output from the left index finger with single-pulse transcranial magnetic stimulation (TMS) when the targeted motor representation was at baseline and when it was selected, nonselected, or irrelevant for action preparation. Here, input was TMS intensity relative to a participant’s resting motor threshold (%RMT), and output was the amplitude of ensuing motor evoked potentials (MEPs). **d**, CS input-output curves derived from pre-task resting state MEP data (n = 39) and fit with a population-based nonlinear mixed effects model using a three-parameter Boltzmann function. Green dots and error bars indicate mean MEP amplitude ± standard deviation at each TMS intensity. Dashed grey lines illustrate parameter estimates.

## Methods

### Ethical Approval

The study was approved by the University of Oregon Institutional Review Board (Study ID 10182017.017) and conformed to the standards set by the *Declaration of Helsinki*, except for registration in a database. All procedures were performed with each participant’s understanding and written consent.

### Participants

Forty-five young, neurologically healthy adults volunteered to participate. All participants self-reported as right-handed and were screened for contraindications to TMS before data collection. Five participants were excluded from data collection due to a high resting motor threshold (≥ 60% maximum stimulator output), making the TMS input-output protocol unfeasible. An additional participant was excluded after data collection due to noncompliance with task instructions. The remaining 39 participants completed the entire study and were available for all statistical analyses (20 females, 19 males; mean age 20.3 yrs., range 18 to 30 yrs.).

### Task protocol

Goal-directed action preparation was assessed with an instructed-delay two-choice reaction time task (Figure 1b). Participants were seated comfortably in front of a computer monitor (59 Hz refresh rate, ∼60 cm viewing distance) with their hands palm-down and shoulder-width apart on a table surface. The task was designed using Psychtoolbox-3 integrated with MATLAB (R2019b; The MathWorks) and a custom response board (Makey Makey v.1.2; Joylabz) for accurate timing. Stimuli were synchronized with task equipment through an analog-to-digital data-acquisition module (PCI-6620; National Instruments) and verified during later offline analyses using photodiode recordings. The default task display consisted of a blank grey background at trial onset. A fixation cue (centered white rectangle, 20 x 20 pixels) appeared 0.3 s after trial onset. A warning cue appeared 1 to 1.5 s after trial onset (randomly drawn from a uniform distribution) and indicated the forthcoming response (hollow leftward or rightward pointed triangle, 140 x 150 pixels). An imperative stimulus (triangle filling white) appeared 0.9 s after the warning cue and lasted until a button press was recorded or 0.8 s elapsed. Each trial was followed by an intertrial interval (blank grey background) after the imperative stimulus and lasted until 7.6 s had elapsed from the trial onset.

The experiment was split into between-hand and within-hand choice tasks (Figure 1c). Both tasks were performed in the same session with a counterbalanced order across participants. Participants were provided a short break in between tasks. Left and right imperative stimuli required button presses from the left and right index fingers during the between-hand choice task and button presses from the right index and pinky fingers during the within-hand task. Participants were instructed to relax during the fixation period, to prepare but not execute the forthcoming response during the cue period, and to execute the cued response as fast as possible when the imperative stimulus was presented. A subset of no-go catch trials (i.e., no imperative stimulus; 8% of trials) was included to discourage participants from responding prematurely in anticipation of the imperative stimulus. For each task, there was a total of 238 trials evenly distributed across left and right imperative stimuli and split across 4 blocks alternating between 59 and 60 trials. The entire experiment lasted 2.5 hours on average.

### Electromyography

Electrophysiological muscle activity was assessed with surface electromyography (EMG) recorded using bipolar Ag-AgCl surface electrodes (Delsys Incorporated). EMG was collected in a copper Faraday cage to minimize electrical noise in EMG recordings. Electrodes were placed on the left and right FDI and the right *abductor digiti minimi* (ADM) hand muscles, which corresponded to the primary agonist muscles for between-hand_left_ (left FDI), between-hand_right_ and within-hand_left_ (right FDI), and within-hand_right_ (right ADM) choice alternatives. An additional EMG electrode was positioned over the C3 vertebra of the neck to record TMS artifacts, and a shared ground electrode was positioned on the left ulnar styloid process. EMG activity was amplified (×1000), bandpass filtered (20 – 450 Hz), baseline-corrected to 0 mV, and sampled at 5000 Hz with a Bagnoli-8 amplifier (Delsys Incorporated) connected to a BNC-2090A terminal block (National Instruments). EMG data were recorded and monitored on each trial for 4 s from onset using the MATLAB-based VETA toolbox (Jackson & Greenhouse, 2019) and stored for later offline analyses.

### Transcranial Magnetic Stimulation

Physiological changes in CS input-output during goal-directed action preparation were assessed with single-pulse TMS delivered using a 70 mm figure-of-eight coil connected to a MagStim 200^2^ stimulator (MagStim Ltd.). The coil was oriented to induce a posterior-to-anterior flowing current in the underlying cortical tissue. The optimal coil position (‘motor hotspot’) over right M1 for eliciting a motor evoked potential (MEP) in the left FDI was then assessed and marked on the scalp. The resting motor threshold (RMT) of left FDI was defined as the minimum stimulator output required to elicit an MEP amplitude of at least 0.05 mV at rest using a maximum-likelihood parameter estimation by sequential testing strategy (Silbert *et al*., 2013). TMS was delivered across a range of stimulation intensities (90%, 110%, 130%, 150%, and 170%; 14 stimuli per intensity) relative to a participant’s RMT (mean 43.4 ± 7.0 % maximum stimulator output) to derive CS input-output curves, with MEP peak-to-peak amplitude used as an index of CS output (Bestmann & Duque, 2016). The TMS input-output protocol was performed at baseline and delay time points in both the between-hand and within-hand tasks. Baseline TMS coincided with the onset of the fixation cue (i.e., end of intertrial interval) to evaluate resting CS output within a task context but before any trial-relevant information was provided (Bakken *et al*., 2024). Delay TMS was delivered 100 ms before the imperative stimulus (i.e., 800 ms into the cue period) to evaluate CS output during action preparation and, in related tasks, corresponds to a time point of maximal MEP modulation before transition to action execution (Lebon *et al*., 2016). The above task design enabled assessment of CS output to the left FDI muscle during baseline and during action preparation when the muscle was selected (i.e., the agonist for prepared action), nonselected, or task-irrelevant (Figure 1c). Nonselected refers to the context in which the left index finger was a choice alternative in the task but was not cued to respond on a given trial, while task-irrelevant refers to the context in which the left index finger was never a choice alternative in the task. Here, we conceptualize the left FDI as task-relevant in both selected and nonselected contexts since CS output is modulated differently when an effector is part of a task set than when it is task-irrelevant entirely (Greenhouse *et al*., 2015).

### Dependent measures

Data were preprocessed using the VETA toolbox in two stages. In the first stage, EMG bursts, MEPs, and TMS artifacts were marked with the automated detection *findEMG.m* function. In the second stage, data was manually reviewed using the *visualizeEMG.m* function to ensure the accuracy of EMG measures. Here, EMG data contaminated by artifacts (e.g., electrical noise, pre-TMS muscle activity) were rejected, and EMG burst onset timings were adjusted when required. Dependent measures were then extracted with custom scripts in MATLAB. Behavioral measures were analyzed from nonstimulated and stimulated trials separately to account for the potential influence of TMS on motor output (Foltys *et al*., 2001). Go success was calculated as the percentage of go trials where a correct response was made within 0.8 s from imperative stimulus onset. Catch trial success was calculated as the percentage of no-go trials where the cued response was withheld correctly. Trials with button press RTs less than 100 ms or EMG RTs less than 50 ms were then marked as premature responses and excluded from subsequent analyses (*M* = 5.8 % of nonstimulated and 13.7% of stimulated trials, respectively). There was a positive correlation between button press and EMG RT for nonstimulated (*r* = 0.68, *P* < 0.001) and stimulated trials (*r* = 0.54, *P* < 0.001), as such, we report only EMG-based RTs for subsequent analyses since it is free from electromechanical delays.

Dependent measures from stimulated trials during the between-hand and within-hand choice tasks included pre-trigger root mean square EMG and MEP amplitudes. In addition to manually reviewing with the VETA toolbox, remaining TMS trials were excluded when average pre-trigger root mean square EMG activity exceeded 30 µV in a -30 to -5 ms pre-TMS window or when outlier MEP amplitudes were identified, defined as values more than three times the scaled median absolute deviation from the median. A mean of 11.4 % of stimulated trials were removed during preprocessing. The mean pre-TMS root mean square EMG activity of the left FDI was then calculated from the remaining trials for each preparation context, collapsed across stimulation intensities. CS output of the left FDI was calculated as the mean MEP peak-to-peak amplitude for each preparation context and stimulation intensity separately.

### Statistical analyses

Statistical analyses were performed using R software (Version 4.3.0). The normality of data and model-averaged residual plots were visually inspected where appropriate. Logarithmic transformations were applied to non-normally distributed data, after which model-averaged residual plots were re-evaluated for normality before hypothesis testing. The alpha level was set to 0.05 for statistical significance. P-values for pairwise post-hoc comparisons were obtained using estimated marginal means for mixed-effects models and adjusted for multiple comparisons using the Benjamini-Hochberg method to control the false discovery rate at 5%. In-text data are presented as nontransformed mean ± standard deviation unless otherwise specified.

We employ a population-based nonlinear mixed-effects model to determine the influence of action preparation context on the CS input-output relationship. A modified three-parameter Boltzmann sigmoid function (Heusser *et al*., 2021) was used to model the nonlinear sigmoidal input-output relationship between TMS intensity and MEP amplitude (Kukke *et al*., 2014):

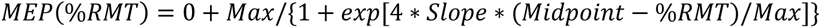

where MEP(%RMT) is the predicted MEP peak-to-peak amplitude at a given TMS intensity, calculated as a function of Max (upper plateau), Midpoint (inflection point), and Slope (maximum rate of change) input-output parameter estimates. Here, the Midpoint and Slope parameters indicate additive and multiplicative computations, respectively (Figure 1d). The lower plateau of CS input-output was constrained to 0 mV for greater biological plausibility (i.e., MEP amplitude cannot be negative and should always be 0 mV at a 0% TMS intensity) and provided a better fit of the data compared to an unconstrained model (constrained AIC: 1307.5, unconstrained AIC: 1317.0).

Initial parameter estimates for the nonlinear mixed-effects models were derived from pre-task resting state data to ensure independence between model optimization and statistical analyses. Specifically, initial estimates were set to mean MEP amplitude at 170% RMT for Max, median stimulation intensity for Midpoint (i.e., 130% RMT), and the linear gradient between 110% and 130% RMT for Slope based on visual inspection of averaged MEP amplitude data. We then applied nonlinear least squares modeling (*nls* function in R) with these initial parameter estimates to obtain corresponding optimal parameter estimates of 2.34 mV for Max, 125.6 %RMT for Midpoint, and 0.055 mV/%RMT for Slope with an iterative numerical integration procedure. These estimates were subsequently applied to a model of in-task TMS data (*nlme* function in R). Max, Midpoint, and Slope parameters were estimated from this random intercept model to account for participant variability. The model was then updated with the fixed effect of Context (baseline, irrelevant, nonselected, selected) to examine the influence of action preparation context on CS input-output. Updated models with the fixed effect of Task Order (within-hand first, between-hand first) or Sex (male, female) were also included to determine any effects of task order or participant sex on CS input-output parameters.

A linear mixed-effects model with a fixed effect of Context (baseline, irrelevant, nonselected, selected) was used to confirm that modulation of CS input-output was not attributable to shifts in pre-TMS root mean square EMG activity. An additional analysis of raw MEPs was included to determine the prevalence of preparatory suppression, defined by a reduction in MEP amplitude relative to baseline (Duque & Ivry, 2009), across the CS input-output spectrum. Here, we used paired Wilcoxon-signed rank tests to compare the mean MEP amplitude for a given preparation context to the baseline context for each suprathreshold stimulation separately.

Go and catch trial success rates for nonstimulated and stimulated trials were analyzed separately using an omnibus Friedman test with the factor of Response (within-hand_left_, within-hand_right_, between-hand_left_, between-hand_right_) to verify task performance was high and consistent across response alternatives. Nonstimulated and stimulated trial EMG RT was analyzed using a linear mixed-effects model with fixed effects of Task (within-hand, between-hand) and Side (left, right) and random effect of participant specificity to verify participants were responding fast and to determine any differences between tasks and response alternatives.

We then estimated a Gain Index *post priori* for each participant to capture interindividual differences in CS input-output Slopes and evaluate whether observed context-dependent gain adjustments correlated with task behavior. Direct parameter estimations could not be used from the population-based model since variance is shared across participants and, thus, interindividual differences between contexts derived from the population-based model are not meaningful. As such, we estimated the Gain Index by 1) extracting the estimated Midpoint for each context and participant separately, 2) determining the two neighboring input TMS intensities (e.g., 110% and 130% RMT if the estimated Midpoint was within these bounds), 3) estimating the linear gradient (slope) between the mean MEP amplitudes from their individual data corresponding to the selected input TMS intensities, and 4) calculating the estimated difference between task-relevant contexts (average of selected and nonselected contexts) and task-irrelevant contexts, normalized to baseline, i.e. (slope_relevant_ - slope_irrelevant_)/slope_baseline_. Thus, a greater Gain Index reflects a larger difference in the Slope of CS input-output between task-relevant and task-irrelevant preparation contexts. The association between mean EMG RT across response options and the Gain Index was then tested using Pearson’s correlation for nonstimulated and stimulated trials separately. The average number of trials available for EMG RT analyses were comparable across response options for both nonstimulated (between-hand_left_ 10.6 ± 1.8; between-hand_right_ 11.7 ± 0.7; within-hand_left_ 11.8 ± 0.5; within-hand_right_ 11.8 ± 1.0) and stimulated (between-hand_left_ 91.6 ± 13.2; between-hand_right_ 92.3 ± 7.0; within-hand_left_ 93.7 ± 7.4; within-hand_right_ 93.8 ± 6.8) trials.

## Results

CS input-output parameters changed during goal-directed action preparation in a context-dependent manner (Figure 2a). The Context model provided a better fit of the data than the Null model (Context AIC: 1264.3, Null AIC: 1312.9). Importantly, there was no effect of Context on pre-TMS muscle activity (*F*_3,144_ = 0.58, *P* = 0.629), which was below 10 µV in all contexts (baseline 5.5 ± 2.5 µV; irrelevant 5.8 ± 4.0 µV; nonselected 5.5 ± 2.7 µV; selected 5.8 ± 2.5 µV). Thus, participants’ muscles rested during action preparation, and differences in suprathreshold alpha motoneuron depolarization did not explain the modulation of CS input-output (Devanne *et al*., 1997).

**Figure 2:**
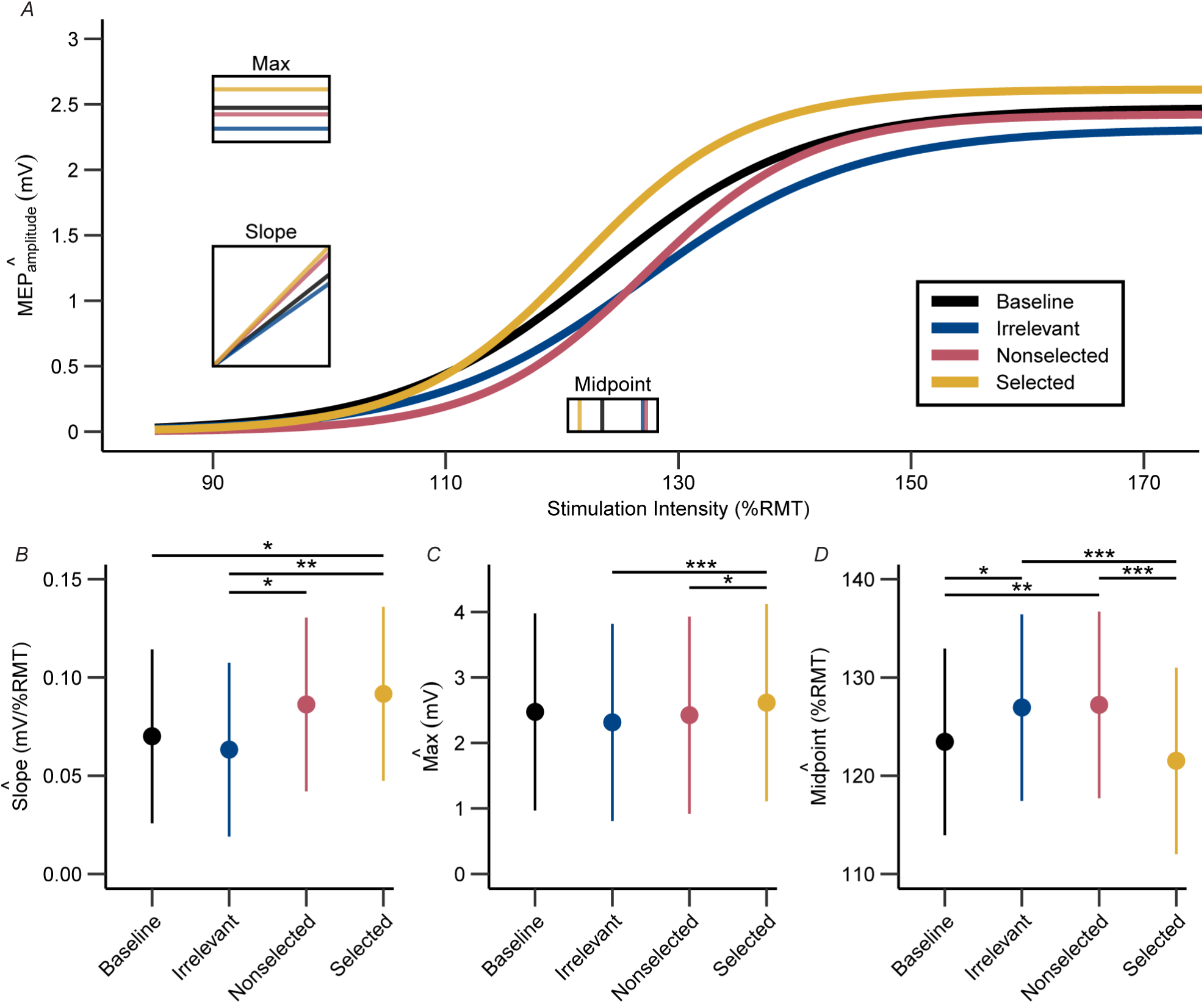
Modulation of CS input-output across goal-directed action preparation contexts. **a**, CS input-output curves across action preparation contexts. Insets indicate the model-derived Max, Slope, and Midpoint parameters used to generate the curves. **b**, Estimated Slope was greater in the selected context compared to baseline and irrelevant, greater in the nonselected context than irrelevant, with no significant differences between baseline and irrelevant or between nonselected and selected contexts. **c**, Estimated Max was greater in the selected context compared to the irrelevant and nonselected contexts, which did not differ from each other, with no differences between baseline and any other contexts. **d**, Estimated Midpoint was greater in the irrelevant and nonselected contexts compared to baseline and the selected context, with no significant differences between nonselected and irrelevant or between baseline and selected contexts. Data ranges represent mean ± standard deviation of the model-derived parameters in panels b – d, where the standard error reflects the precision of the estimated coefficients. Asterisks indicate statistical significance, where *P* is * < 0.05, ** < 0.01, *** < 0.001; n = 39 in all panels.

### CS gain increases during response preparation for task-relevant representations

The Slope of CS input-output differed significantly across Context (*F*_3,730_ = 4.26, *P* = 0.00540). Estimated Slope was greater in the selected context compared to baseline (*F*_1,730_ = 5.42, *P* = 0.0404) and the irrelevant context (*F*_1,730_ = 9.76, *P* = 0.0111). Estimated Slope in the nonselected context was also greater than the irrelevant context (*F*_1,730_ = 5.87, *P* = 0.0404) but not baseline (*F*_1,730_ = 2.79, *P* = 0.143). There were no significant differences between baseline and irrelevant (*F*_1,730_ = 0.95, *P* = 0.397) as well as between nonselected and selected contexts (*F*_1,730_ = 0.23, *P* = 0.628). Thus, context-dependent multiplicative modulation of CS input-output was explained by an increase of Slope when the stimulated motor representation was selected or nonselected for forthcoming action (i.e., task-relevant) compared to baseline or when it was task-irrelevant (Figure 2b). Such nonlinear increases in sensitivity to excitatory inputs are associated with response potentiation across sensory systems (Wood *et al*., 2017). Multiplicative gain modulation may be important for maximizing signal-to-noise and, computationally, can extend the dynamic range of neuronal operations (Mitchell & Silver, 2003; Shine *et al*., 2018). Of note is that increased Slope was observed in both the selected and nonselected preparation contexts, indicating a general role of gain modulation in potentiating task-relevant motor representations rather than promoting effector-specific action selection. Overall, gain modulation may be pertinent for establishing the pool of candidate actions.

The Max of CS input-output also differed significantly across Context (*F*_3,730_ = 24.04, *P* < 0.001). Estimated Max was greater in the selected context compared to irrelevant (*F*_1,730_ = 14.62, *P* < 0.001) and nonselected (*F*_1,730_ = 6.44, *P* = 0.0341) contexts, which did not differ from each other (*F*_1,730_ = 1.89, *P* = 0.204). There were no differences between baseline and irrelevant (*F*_1,730_ = 3.85, *P* = 0.0978), nonselected (*F*_1,730_ = 0.40, *P* = 0.528), or selected (*F*_1,730_ = 3.41, *P* = 0.0978) contexts. Thus, CS output maximum was greater during selected than nonselected and irrelevant contexts, but not baseline (Figure 2c). This pattern differed from that observed for the Slope and indicates the dynamic range of excitability in the CS pathway is influenced by more than gain adjustments alone.

### Additive suppression of CS input-output in nonselected and task-irrelevant preparatory contexts

The Midpoint of CS input-output differed significantly across Context (*F*_3,730_ = 11.48, *P* < 0.001). Estimated Midpoint was greater in the irrelevant context than baseline (*F*_1,730_ = 7.00, *P* = 0.0125) and the selected context (*F*_1,730_ = 18.54, *P* < 0.001). Estimated Midpoint in the nonselected context was also greater than baseline (*F*_1,730_ = 9.65, *P* = 0.00393) and the selected context (*F*_1,730_ = 25.32, *P* < 0.001). There was no significant difference between nonselected and irrelevant (*F*_1,730_ = 0.05, *P* = 0.828) and between baseline and selected contexts (*F*_1,730_ = 2.50, *P* = 0.137). Thus, context-dependent modulation of CS input-output was explained by an additive rightward shift of the Midpoint when the stimulated motor representation was task-irrelevant or nonselected for the forthcoming action compared to when it was selected (Figure 2d). The additive increase of the Midpoint during nonselected and irrelevant contexts was also greater than baseline, consistent with inhibition of CS input-output relative to intertrial rest (Derosiere & Duque, 2020). This nonspecific and apparently uniform suppression of irrelevant and nonselected motor representations may reflect widespread noise reduction within the CS pathway (Greenhouse *et al*., 2015). These findings extend previous studies of preparatory inhibition and are consistent with a mechanism for enhancing the contrast of selected motor representations through the suppression of background activity.

### Preparatory suppression during goal-directed action preparation

Preparatory suppression, expressed as a reduced MEP amplitude relative to baseline, was observed during the irrelevant context across all stimulation intensities (110: *W* = 615, *P* = 0.00374; 130: *W* = 573, *P* = 0.0293; 150: *W* = 608, *P* = 0.00546; 170: *W* = 616, *P* = 0.00354), only at the 110 %RMT stimulation intensity during the nonselected context (110: *W* = 644, *P* < 0.001; 130: *W* = 501, *P* = 0.372; 150: *W* = 411, *P* = 1.00; 170: *W* = 446, *P* = 1.00), but at no stimulation intensity during the selected context (110: *W* = 393, *P* = 1.00; 130: *W* = 238, *P* = 0.100; 150: *W* = 271, *P* = 0.296; 170: *W* = 280, *P* = 0.382). Thus, preparatory suppression was observed across the entire input spectrum in the irrelevant context, only at the lower end of the input spectrum in the nonselected context and at no part of the input spectrum in the selected context (Figure 3a).

**Figure 3:**
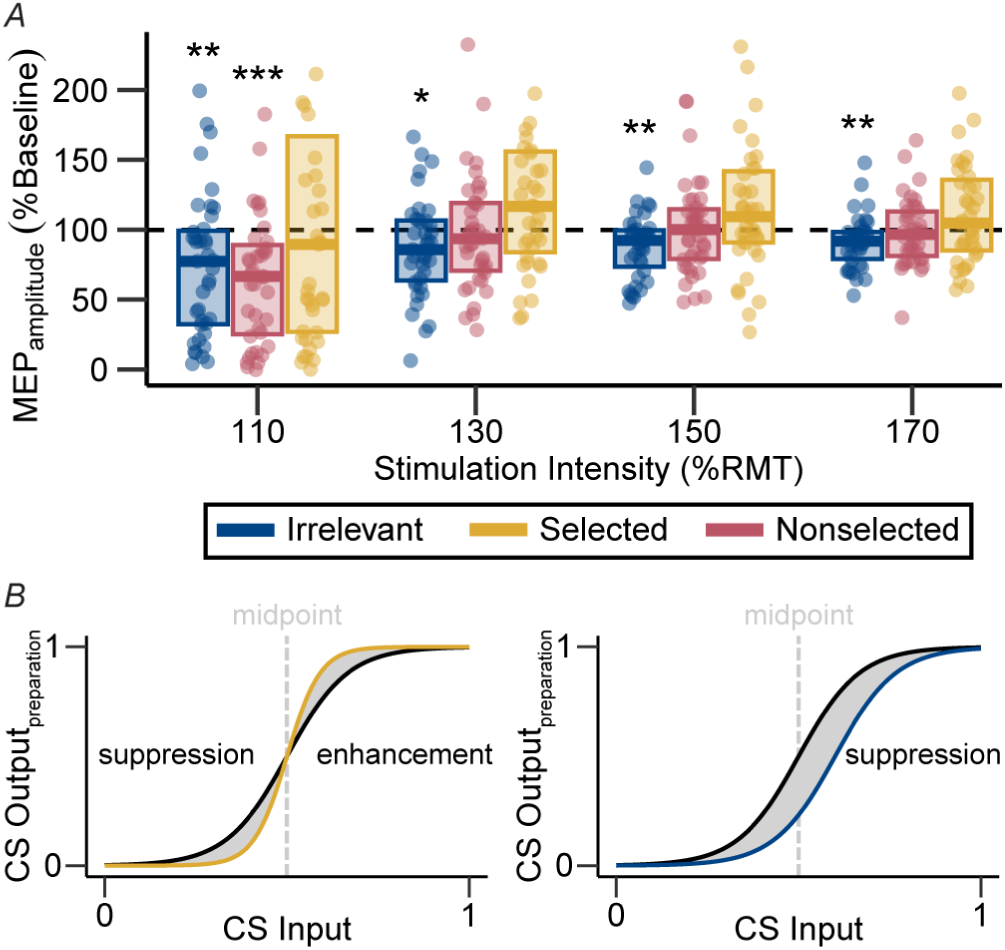
Preparatory suppression during goal-directed action preparation. **a**, Preparatory suppression, expressed as a reduced MEP amplitude relative to baseline, was observed during the irrelevant context across all stimulation intensities, only at the 110 %RMT stimulation intensity during the nonselected context, and at no stimulation intensity during the selected context. Crossbar boxes indicate sample median, 25^th^, and 75^th^ percentiles. The dashed horizontal line indicates baseline MEP amplitude. Asterisks indicate statistical significance, where *P* is * < 0.05, ** < 0.01, *** < 0.001; n = 39. **b**, Conceptual overview of how CS input-output modulation can reflect preparatory suppression or enhancement depending on CS input and modulation. Here, assuming fixed alternative parameters, a multiplicative increase would be captured as preparatory suppression or enhancement at input intensities below or above the midpoint, respectively. In contrast, an additive increase (i.e., rightward shift) would be captured as preparatory suppression irrespective of the input intensity relative to the midpoint.

### Gain modulation of CS input-output across preparatory contexts is associated with reaction time

Participants responded accurately and quickly during the behavioral tasks. There were no significant differences in accuracy across the response options during either go (nonstimulated: *X*^2^ = 4.82, *P* = 0.186; stimulated: *X*^2^ = 5.10, *P* = 0.164) or catch (nonstimulated: *X* ^3^ = 3.00, *P* = 0.392; stimulated: *X*^2^ = 3.12, *P* = 0.373) trials, with a high average trial success indicating that participants performed the tasks correctly (Figure 4a). For EMG RT during nonstimulated trials, there was a significant fixed effect of Task (*F*_1,114_ = 10.18, *P* = 0.00183), with faster responses in the within-hand compared to the between-hand task. There was no fixed effect of Side (*F*_1,114_ = 0.17, *P* = 0.684) or a Task × Side interaction (*F*_1,114_ = 2.88, *P* = 0.0924). These EMG RT results were consistent during stimulated trials, whereby responses were faster in the within-hand compared to between-hand task (*F*_1,114_ = 6.56, *P* = 0.0117), with no fixed effect of Side (*F*_1,114_ = 0.15, *P* = 0.699) or a Task × Side interaction (*F*_1,114_ = 0.63, *P* = 0.430). Thus, responses were faster when the prepared action involved a within-hand choice between the fingers of the dominant hand compared to a between-hand choice. However, RTs remained fast across all response alternatives (Figure 4b). Importantly, RTs were correlated between the two tasks for nonstimulated (*r* = 0.68, *P* < 0.001) and stimulated (*r* = 0.38, *P* < 0.001) trials, indicating individuals differed reliably in their RTs. There was a significant negative Pearson correlation between the Gain Index and EMG RT from nonstimulated (*r* = -0.32, *P* = 0.0449) and stimulated (*r* = -0.39, *P* = 0.0131) trials, whereby participants with a larger gain modulation index had faster RTs (Figure 4c). That is, participants who exhibited greater gain adjustments in the form of slope differences between task-relevant and task-irrelevant behavioral contexts were faster to initiate responses.

**Figure 4:**
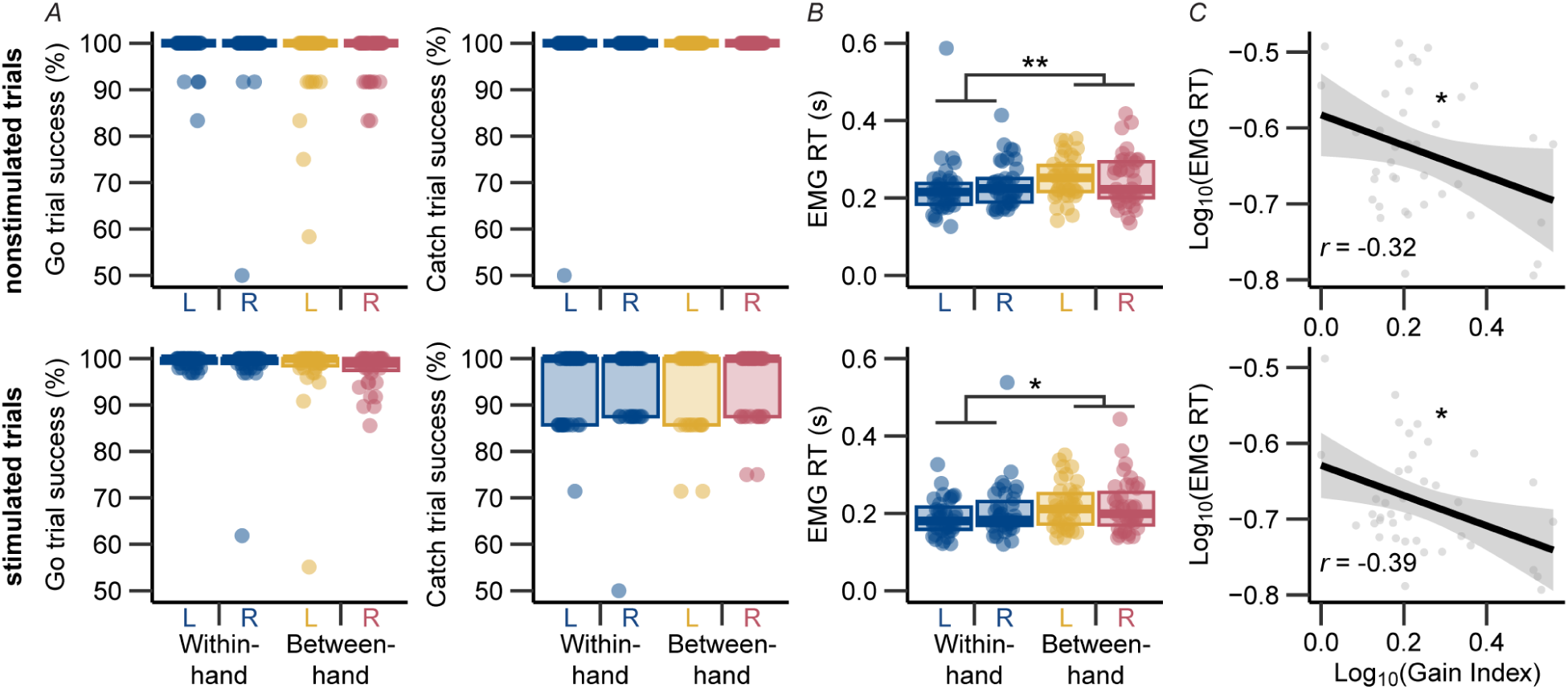
Behavioral performance during nonstimulated (*top*) and stimulated (*bottom*) trials and its relationship to CS input-output gain modulation. **a**, Accuracy during go (left) and catch (right) trials for which there was no significant effect of Response type on either measure. **b**, Response initiation times based on EMG were faster during the within-hand compared to the between-hand task. **c**, There was a significant negative Pearson correlation between Gain Index and EMG RT, whereby participants with greater gain in task-relevant than task-irrelevant action preparation contexts were faster responders. Crossbar boxes indicate sample median, 25^th^, and 75^th^ percentiles. Asterisks indicate statistical significance, where *P* is * < 0.05, ** < 0.01; n = 39 in all panels.

### CS input-output is not affected by participant sex or task order

CS input-output curves derived from the Sex model did not provide a better fit of the data than the null model (Sex AIC: 1319.1, Null AIC: 1312.9). There were no significant effects of Sex on estimated Max (*F*_1,736_ = 0.61, *P* = 0.434), Slope (*F*_1,736_ = 0.27, *P* = 0.605), or Midpoint (*F*_1,736_ = 2.85, *P* = 0.0916) CS input-output parameters. CS input-output curves derived from the Order model also did not fit the data better than the null model (Order AIC: 1320.1). There were no significant effects of Order on estimated Max (*F*_1,736_ = 0.04, *P* = 0.844), Slope (*F*_1,736_ < 0.01, *P* = 0.944), or Midpoint (*F*_1,736_ = 0.02, *P* = 0.876) CS input-output parameters. Thus, neither participant Sex nor task order influenced CS input-output during goal-directed action preparation (Figure 5).

**Figure 5:**
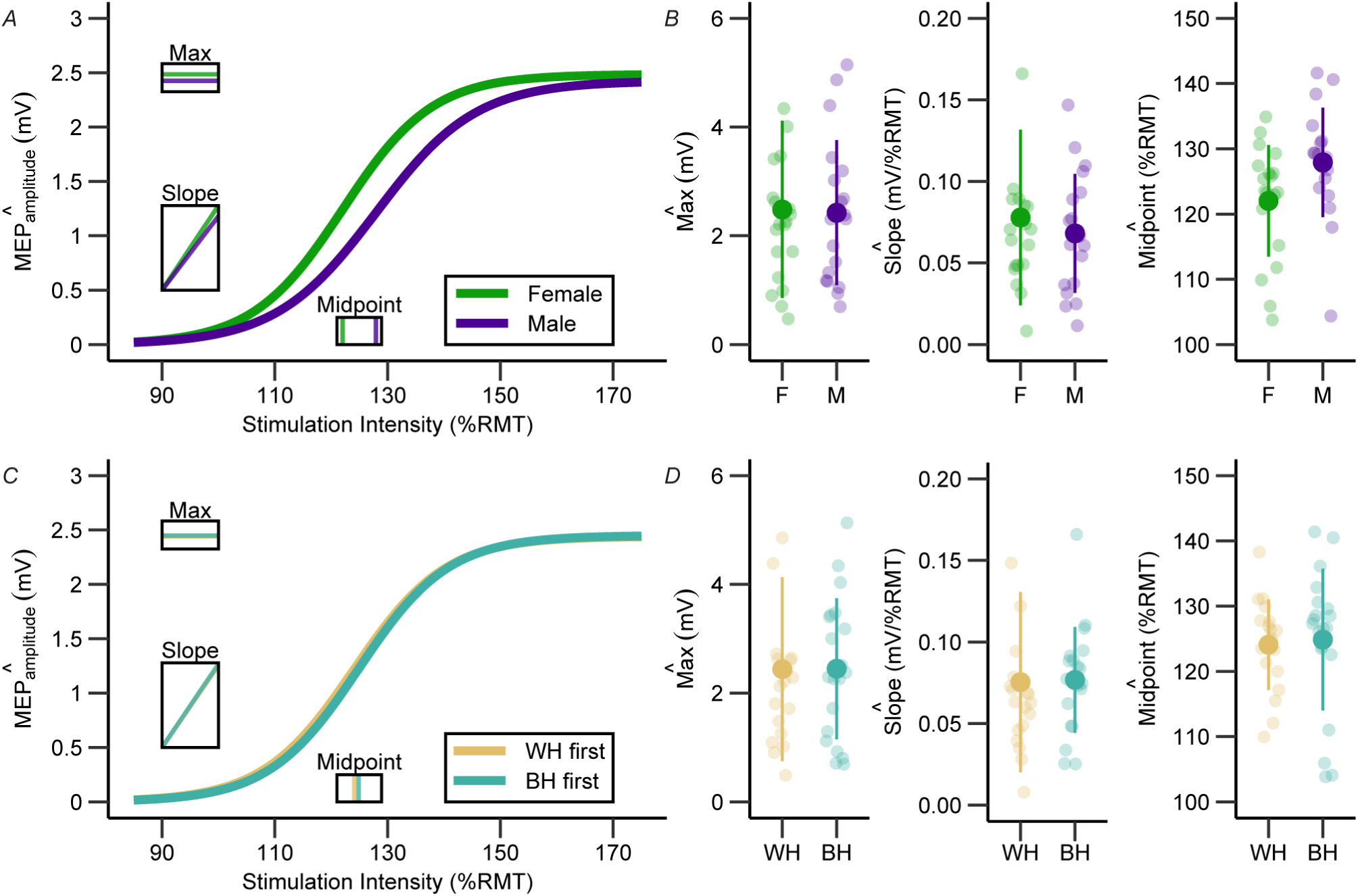
Modulation of CS input-output across participant sex and task order. **a**, CS input-out curves derived from the Sex model. **b**, There were no significant effects of Sex on estimated Max, Slope, or Midpoint CS input-output parameters. **c**, CS input-output curves derived from the Order model. **d**, Task order (WH: within-hand first; BH: between-hand first) had no significant effects on estimated Max, Slope, or Midpoint CS input-output parameters. Insets in subpanels a and c indicate estimated population means for the Max, Slope, and Midpoint parameters. Data ranges in subpanels b and d represent mean ± standard deviation of the model-derived parameters, where the standard error reflects the precision of the estimated coefficients. Individual datapoints indicate extracted participant-wise estimates of Max, Slope, and Midpoint parameters for between-subjects effects. n = 39 in all panels.

## Discussion

The present study provides novel physiological evidence of multiplicative and additive neural computations within the human CS pathway during goal-directed action preparation. Specifically, we showed that the gain of behaviorally relevant motor representations (selected and non-selected) increases against a background of additive suppression of nonselected and task-irrelevant motor representations during goal-directed action preparation. Furthermore, the magnitude of gain modulation was associated with behavioral output, such that stronger modulators were faster responders. Below, we discuss the implications of these findings for understanding state-dependent computations within the human CS pathway during motor control.

### Computational parallels between sensory and motor systems during action preparation

First, the findings indicate canonical computations well-described in sensory cortices (Salinas & Sejnowski, 2001; Mitchell & Silver, 2003; Silver, 2010) generalize to the human motor system. Previous work has established feature-based attention modulates gain within visual cortices (Treue & Trujillo, 1999), and normalization in the human visual system further supports attentional processes by theoretically separating figures from the ground (Bloem & Ling, 2019). Similar gain adjustments have been observed in the auditory cortex to increase spectrotemporal contrast (Rabinowitz *et al*., 2011). Computational models of auditory (Mischler *et al*., 2023) and visual (Polack *et al*., 2013) sensory systems further indicate specific roles of gain for normalization and may mirror the contrast enhancement mechanism we propose operates within the CS pathway during action preparation. Such state-dependent gain adjustments for contrast enhancement could facilitate the transmission of descending signals along the CS pathway, similar to models of basal ganglia output for action selection (Nambu *et al*., 2002). Importantly, while sensory and motor systems differ in their representational content and anatomy, they share critical functional similarities for guiding behavior (Hatsopoulos & Suminski, 2011). Therefore, contrast enhancement via simultaneously increased gain of behaviorally relevant and suppression of background neural representations may be a canonical motif for sculpting cortical circuit output.

The current findings also align with recurrent neural network models of the non-human primate motor cortex that indicate a specific role of gain modulation in moderating movement features such as speed (Stroud *et al*., 2018). Such models are typically based on electrophysiological recordings from a neuronal subpopulation in a target region acquired without experimental control over the input strength to the efferent motor pathway. Here, by controlling the input intensity and tailoring it to each participant, we provide corroborative physiological evidence for these models by demonstrating that gain modulation along the human CS pathway during a preparatory state facilitates the execution of subsequent actions. Based on our data, we propose that increased gain of task-relevant motor representations occurs against a backdrop of suppressed nonselected and irrelevant motor representations to enhance the contrast between selected and bystander motor representations (Greenhouse *et al*., 2015). We theorize this extension to existing models further promotes the initiation of prepared responses by increasing signal-to-noise within the CS pathway via divisive normalization (Figure 6). Our data can also inform computational models of the motor cortex by providing useful information about the dynamic range of physiological outputs and behavioral relationships in a human model. Translation of the context-dependent approach to other model systems can help evaluate whether CS input-output adjustments during an output-null (preparatory) motor state hasten the transition to an output-potent (execution) motor state (Hannah *et al*., 2018; Churchland & Shenoy, 2024).

**Figure 6:**
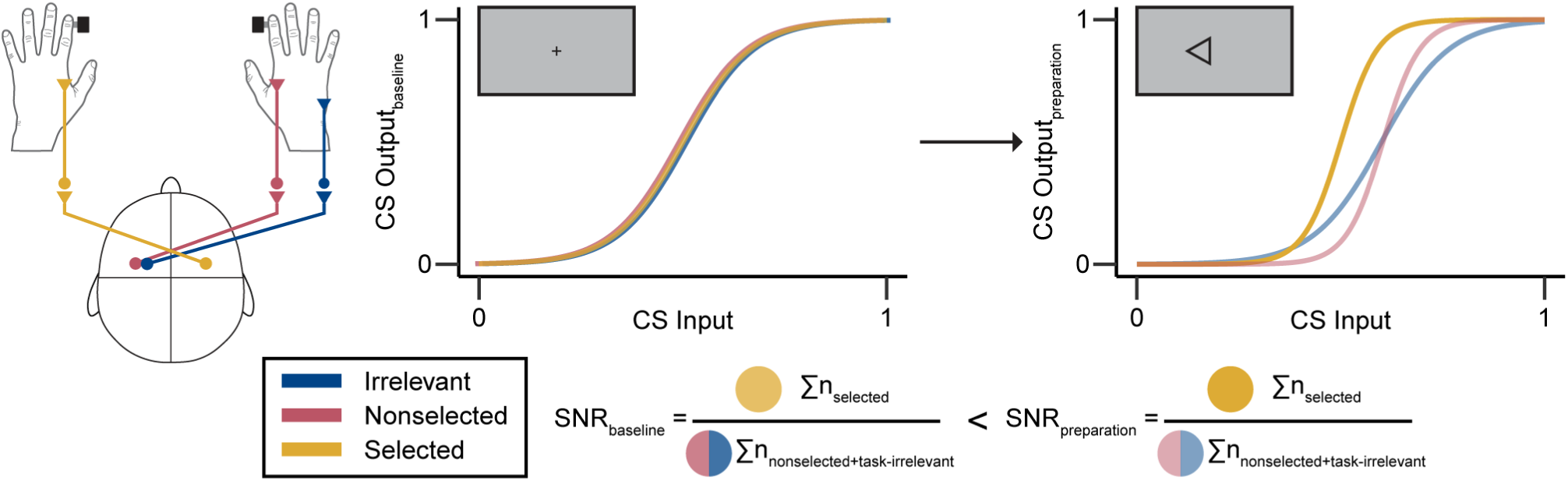
Theorized role of divisive normalization during goal-directed action preparation. CS input-output during action preparation depends upon task contexts. Schematized CS input-output plots represent the multiplicative increase in the selected and nonselected contexts and the additive increase in nonselected and task-irrelevant contexts. Speculatively, complementary multiplicative and additive changes can enhance the signal-to-noise ratio (SNR) between selected and bystander motor representations through divisive normalization to improve the fidelity of descending CS commands during action preparation.

### Context-dependent computations with common coding frameworks

Context-dependent computations during goal-directed action preparation may fit within cognitive ‘common coding’ frameworks. These frameworks posit that human action control tasks are supported by internal representations (‘event files’) of stimulus-response relationships and their sensory effects (Hommel, 2009). Accordingly, action preparation entails retrieving an event file that triggers integrated neuronal processing across nonmotor and motor brain regions to prime action execution (Beste *et al*., 2023). We observed neurophysiological evidence of corticospinal input-output computations recapitulated from sensory systems during action preparation, which may suggest a perception-action common code between motor and nonmotor systems. The current results, conceptualized within a common code framework, indicate that event files may have widespread motoric effects on not only the CS pathway to effectors involved directly in stimulus-response bindings but also suppression of task-irrelevant CS effector representations. In a somewhat homologous manner, modulation of sensory feedback during action control occurs at both spinal and supraspinal levels and can include a mixture of gain and suppression computations (Azim & Seki, 2019). Binding the motor command to anticipated perceptual feedback may occur via common gain modulation and suppression processes, with up-or-down regulation of neural pathways contingent upon their behavioral relevance (Frings *et al*., 2020). Event file bindings are likely strong in the instructed-delay response task we employed; however, it is important to note that we did not set out to test event files explicitly and did not manipulate binding versus retrieval processes. Nevertheless, our findings raise the interesting possibility that gain increases in CS output during goal-directed action preparation are reciprocal to gain decreases in cutaneous feedback (Seki & Fetz, 2012) but coupled to gain increases in proprioceptive feedback during action execution (Confais *et al*., 2017). Future work can investigate whether such changes arise from a common descending command.

### Reevaluating context-dependent preparatory suppression across the CS input spectrum

An interesting observation from the current study was the lack of MEP suppression in the selected context. Previous TMS studies have observed *preparatory suppression* (i.e., reduced MEP amplitude below baseline during action preparation) across selected, nonselected, and task-irrelevant contexts, albeit variably (Bundt & Huster, 2024). Our finding of enhanced CS gain in task-relevant contexts may appear in conflict with inhibitory accounts (Ibáñez *et al*., 2020). However, previous studies often used a TMS intensity of ≤ 120 %RMT and, thus, only captured CS output at the lower end of the input spectrum. By using a range of TMS intensities, we could model the entire CS input-output function, providing a more complete picture of the underlying processes. Mathematically, both additive and multiplicative changes in a sigmoidal input-output function would have complex effects on whether context-dependent modulation is captured as suppression or enhancement when input intensity is below or above the Midpoint (Figure 3b). We speculate that previous evidence of preparatory suppression of task-relevant motor representations may result from TMS intensities below the CS input-output Midpoint. Importantly, TMS may not mimic endogenous input to the CS pathway when transitioning from action preparation to execution, which remains open for further investigation.

### Neural basis of context-dependent gain adjustments along the CS pathway

Simultaneous gain adjustments and the suppression of background activity along the entire CS pathway may rely on cooperative cortical, subcortical, and spinal mechanisms. For example, subcortical-cortical projections from the thalamus to the prefrontal cortex selectively influence signal (figure) and noise (ground) to guide cued behavior in the context of uncertainty via two distinct intracortical mechanisms (Mukherjee *et al*., 2021). Similar motifs may exist in thalamic projections to M1 to modulate CS output (Halassa & Sherman, 2019). According to this framework, both increased gain of selected motor representations and suppressed background activity would arise from a modulatory influence of intracortical inhibitory circuits (Ferguson & Cardin, 2020). Alternatively, inhibition of background activity may reflect the engagement of CS projections to spinal inhibitory neurons to suppress muscles that, if activated, would interfere with the execution of the prepared action (Griffin & Strick, 2020).

Computations may also be mediated directly by spinal motoneurons, as MEPs measured with surface EMG exhibit sigmoidal input-output properties, but individual spinal motor unit discharge rates exhibit linear input-output properties (Devanne *et al*., 1997). Summations across pools of spinal motoneurons operating around the subliminal fringe may contribute to nonlinear transformations at the population level (Di Lazzaro *et al*., 1999), producing the sigmoidal shape modeled in our data. This account indicates that population-level activity of spinal motoneurons operating more closely to an activation threshold during action preparation could be responsible for the context-dependent gain changes we observed, independent of cortical mechanisms. However, evidence of altered corticospinal input-output properties, such as elevated gain in Parkinson’s disease (Bologna *et al*., 2018) and reduced maximum MEP output after stroke (Buetefisch *et al*., 2018), underscores the importance of cortico-subcortical mechanisms. Therefore, it is likely that the observed CS input-output computations depend on cooperative cortical and spinal mechanisms rather than a single stage in the pathway.

Intra-hemispheric and inter-hemispheric mechanisms are engaged during action preparation (Derosiere & Duque, 2020) and undoubtedly contribute to the observed context-dependent computations. Regarding intra-hemispheric mechanisms, context-dependent computations may reflect distinct contributions to MEPs at varying TMS intensities. TMS-induced MEPs result from summated volleys of indirect (polysynaptic) and direct activation of pyramidal tract neurons along the CS pathway (Di Lazzaro *et al*., 2012). The recruitment pattern is partially intensity-dependent, such that near-threshold TMS tends to evoke early indirect activation while increasing the intensity evokes later indirect activation and, ultimately, direct activation at high intensities (Di Lazzaro *et al*., 2018). Thus, context-dependent CS input-output modulation may reflect changes in the relationship of indirect to direct waves due to the anatomical sources of MEPs (output) shifting with intensity (input). However, such changes could only be attributed to intrinsic differences between contexts, given the matched TMS intensities across the tasks. Inter-hemispheric projections via the corpus callosum are also engaged during action preparation (Hinder *et al*., 2018; Wadsley *et al*., 2023). Callosal projections in the mammalian brain are anatomically organized such that reciprocal projections to inhibitory interneurons and pyramidal tract neurons could support an interplay between excitatory and inhibitory processes (O’Hashi *et al*., 2018). As such, contrast enhancement via simultaneous gain adjustments and ‘crossed’ surround inhibition may be an emergent functional property of inter-hemispheric connectivity in the motor system (for a detailed review, see Carson, 2020). The current study supports this integrative role of inter-hemispheric mechanisms given that nonselected and task-irrelevant computations both exhibited additive suppression but differed in gain, despite both being assessed in the opposite hemisphere of the selected effector. Ultimately, action preparation likely requires integrated processing across intra-hemispheric and inter-hemispheric mechanisms to provide for truly context-dependent computations.

### Limitations & Future Directions

The present study has some limitations. Using motor-evoked potentials (MEPs) elicited by TMS provides an accurate assay of CS output. However, when probed with single-pulse TMS, MEP modulation can arise from various mechanisms along the entire CS pathway, including intracortical, transcortical, subcortical, and spinal mechanisms (Duque *et al*., 2017), which may have unique roles in action preparation (Derosiere & Duque, 2020). Thus, while we demonstrate that various computations manifest in CS output, we cannot disentangle the level of the motor system in which they arise. Future experiments will be able to determine at which level within the CS pathway these computations occur by leveraging multimodal approaches, such as paired-pulse TMS, to characterize better the multifaceted neuronal response to noninvasive brain stimulation and by studying clinical populations known to have CS input-output irregularities. Similarly, future experiments can record MEPs from task-irrelevant muscles within the same hemisphere as the selected effector to better determine the spatial specificity of CS input-output computations. This approach could also help disentangle the role of intra-hemispheric and inter-hemispheric mechanisms during action preparation.

Another limitation relates to in-task TMS only being delivered at baseline or a fixed time point late during action preparation. Limiting the temporal sampling was an important methodological choice to collect a reliable number of trials for accurate CS input-output modeling across various contexts. However, the time course of the observed computations during the transition from action preparation to execution remains unclear. Future experiments may employ methods to sample CS input-output effectively to enable a greater temporal resolution during action preparation (e.g., Alavi et al., 2019). Similarly, MEPs were recorded from only one muscle involved in the various response options, which limited comparisons of CS input-output computations with behavior. Future experiments may use advanced TMS techniques to simultaneously evaluate CS input-output in left and right M1 during action preparation (e.g., Vassiliadis et al., 2018).

The current work did not directly test relationships between the common coding framework and motor system anatomy. However, the observed mixture of context-dependent gain modulation and additive suppression may bridge these seemingly distinct concepts. Specifically, subcortical, intracortical, transcortical, or inter-hemispheric modulatory influences on the CS pathway might provide the neural infrastructure for common coding between sensory and motor areas (e.g., Takacs et al., 2021). Disruptions to these mechanisms could impair the perception-action common code for stimulus-response bindings. While this study does not favor one explanation over another, future studies mapping computations to neuroanatomical mechanisms could clarify how event files are instantiated and retrieved during action preparation.

### Conclusion

The present study aimed to test the presence of multiplicative and additive computations in human CS input-output during goal-directed action preparation. We provided physiological evidence that goal-directed action preparation of finger responses entails a mixture of input-output computations within the human motor system. These computations are context-dependent, characterized by gain increases in task-relevant motor representations against a backdrop of suppression in bystander motor representations, potentially facilitating action initiation through contrast enhancement. The recapitulation of canonical neural computations across motor and non-motor neuronal circuits suggests that these processes may generalize throughout the human nervous system.

## Additional information

### Data availability statement

The data and reproducible code that support the findings of this study are openly available in OSF at https://osf.io/837ms

## Acknowledgments

The authors thank Kate Bakken, Mitch Fisher, Charlie Lewkowitz, Hayami Nishio, Michelle Ortman, and Tania Sarabia for their help with data collection and preprocessing. The authors also thank Luca Mazzucato and Richard Ivry for their helpful feedback on this manuscript.

## Author contributions

Conceptualization, C.G.W and I.G.; Methodology, C.G.W, T.N., and I.G.; Formal Analysis, C.G.W. and T.N.; Investigation, C.G.W. and C.H.; Writing – Original Draft, C.G.W. and I.G.; Writing – Reviewing & Editing, C.G.W, T.N., C.H., and I.G.; Visualization, C.G.W.; Supervision, I.G.; Funding acquisition, I.G.

## Competing interests

The authors declare no competing interests.

## Funding

This research was supported by NIH grant NINDS R01NS123115.

## Notes

### Competing Interest Statement

The authors have declared no competing interest.

### Summary of Updates

Minor revisions to discussion. OSF link to repository is now public.

